# The gammaherpesvirus 68 viral cyclin facilitates reactivation by promoting latent gene expression

**DOI:** 10.1101/761700

**Authors:** Brian F Niemeyer, Joy E Gibson, Jennifer N Berger, Lauren M Oko, Eva Medina, Eric T Clambey, Linda F van Dyk

**Author notes:** Address correspondence to: Linda F. van Dyk, University of Colorado School of Medicine, 12800 E. 19th Ave., MS8333, P.O. Box 6511, Aurora, CO, 80045.

## Abstract

Gammaherpesviruses establish life-long infections within their host and have been shown to be the causative agents of devastating malignancies. Chronic infection within the host is mediated through cycles of transcriptionally quiescent stages of latency with periods of reactivation into more active lytic and productive infection. The mechanisms that regulate reactivation from latency remain poorly understood. Previously, we defined a critical role for the viral cyclin in promoting reactivation from latency. Disruption of the viral cyclin had no impact on the frequency of cells containing viral genome during latency, yet it remains unclear whether the viral cyclin influences latently infected cells in a qualitative manner. To define the impact of the viral cyclin on latent gene expression, we utilized a viral cyclin deficient variant expressing a LANA-beta-lactamase fusion protein (LANA::βla), to enumerate both the cellular distribution and frequency of latent gene expression. Disruption of the viral cyclin did not affect the cellular distribution of latently infected cells, but did result in a significant decrease in the frequency of cells that expressed LANA::βla across multiple tissues and in both immunocompetent and immunodeficient hosts. Strikingly, whereas the cyclin-deficient virus had a reactivation defect in bulk culture, sort purified cyclin-deficient LANA::βla expressing cells were fully capable of reactivation. These data emphasize that the γHV68 latent reservoir is comprised of at least two distinct stages of infection characterized by differential latent gene expression, and that a primary function of the viral cyclin is to promote latent gene expression within infected cells in vivo.

**AUTHOR SUMMARY:** Gammaherpesviruses are ubiquitous viruses with oncogenic potential that establish latency for the life of the host. These viruses can emerge from latency through reactivation, a process that is controlled by the immune system. Control of viral latency and reactivation is thought to be critical to prevent γHV-associated disease. This study focuses on a virally-encoded cyclin that is required for reactivation from latency. By characterizing how the viral cyclin influences latent infection in pure cell populations, we find that the viral cyclin has a vital role in promoting viral gene expression during latency. This work provides new insight into the function of a virally encoded cyclin in promoting reactivation from latency.

## Introduction

Gammaherpesviruses (γHV) are a group of lymphotropic viruses within the herpesviridae family, including the human pathogens Epstein-Barr virus (EBV) and Kaposi’s sarcoma-associated herpesvirus (KSHV, HHV-8). Infection with these viruses is known to result in development of a wide range of malignancies including Burkitt’s lymphoma, Kaposi’s sarcoma, nasopharyngeal carcinoma, post-transplant lymphoproliferative disorders, and primary effusion lymphoma [1, 2]. The naturally occurring mouse gammaherpesvirus, γHV68, is closely related to both EBV and KSHV, readily infects laboratory strains of mice, and provides insights into the complex processes of γHV pathogenesis [3, 4].

γHV infection can be characterized by two distinct phases, lytic and latent infection. Lytic infection is a productive form of infection in which the entire suite of viral genes is expressed and the virus actively replicates its genome [5-7]. In this process, new virus is produced and the lytically infected cell dies. Alternatively, the virus may enter a latent state of infection, in which viral gene expression is mostly suppressed and the viral genome is maintained as an episome in the host nucleus [8]. γHV are able to switch from latent to lytic infection through a process known as reactivation [9, 10]. These viruses are able to establish latent infection in many different cell types including dendritic cells, macrophages, and multiple B cell subsets (including memory B cells, plasma cells, B1-a cells, and B1-b B cells) [11-17]. Although several cell types support latent infection, the relative efficiency of these cell types to support reactivation remains unknown. Numerous studies suggest that a primary source of reactivating virus is plasma cells [14, 18-20]. Other studies indicate that in the peritoneal compartment, infected macrophages and/or B1 B cells are major cell types capable of reactivation [12, 13].

Many viral and host factors contribute to the control of latent infection and reactivation. KSHV and γHV68 both encode a conserved viral cyclin (v-cyclin), which is homologous to host D-type cyclins [3, 21, 22]. Although EBV does not encode its own cyclin, it expresses viral genes that upregulate host cyclin D2, fulfilling a similar function to the KSHV and γHV68 v-cyclin [23]. Like the host cyclins, the v-cyclin has the ability to interact with host cyclin-dependent kinases (CDKs) and promote cell cycle progression [24, 25]. Unlike conventional host cyclins, the v-cyclin is resistant to inhibition by CDK inhibitors (CKI) [26]. Recent work by our group showed that one mechanism by which the v-cyclin promotes reactivation is by antagonizing the host CKI p18Ink4c, in a cell intrinsic manner [27, 28].

Although the v-cyclin is required for reactivation from latency, the underlying mechanisms by which it promotes reactivation have yet to be elucidated. Here, we studied how the v-cyclin may influence latent gene expression in vivo, through the use of recombinant γHV68 viruses that encode a fusion of the ORF73/LANA latency-associated gene with β-lactamase, a robust enzymatic reporter gene that can be used to identify individual virally-infected cells [28, 29], referred to as LANA::βla. By comparing wild-type and cyclin-deficient viruses, we were able to quantify the frequency and cellular distribution of LANA::βla gene expression during latency. These studies demonstrate that the v-cyclin has a critical role in promoting expression of LANA::βla at the single-cell level, with no discernable impact on the cellular distribution of infection. Further, we find that the v-cyclin is completely dispensable for reactivation, when reactivation efficiency is tested in LANA::βla expressing cells. The work detailed here serves to further our understanding of how the virus regulates reactivation. We also highlight an emerging trend in the field of virology where latency is not a uniform state of infection. Rather, some latently infected cells are poised for reactivation, while other infected cells appear to be refractory to reactivation.

## Results

### A cycKO virus expressing a fusion between LANA and β-lactamase is equivalent to wild-type virus in LANA::βla expression during lytic infection, but deficient in reactivation

The v-cyclin is required for γHV68 reactivation. Virus lacking v-cyclin, cycKO, is equivalent to wild-type virus in replication and establishment of latency, but is selectively defective in reactivation from latency [30, 31]. Given that some cell types may be more permissive to reactivation from latency than others, we proposed that the cycKO virus may be enriched in, or limited to, a “less permissive” cell type. To address this, we made use of two previously described enzymatically marked viruses, WT.βla and cycKO.βla [28, 29]. These viruses both contain a fusion protein where β-lactamase is fused to the viral LANA (Fig 1A). This can be used to efficiently identify infected cells by flow cytometry using LANA::βla expression as a surrogate indicator of virus infection. Fusion of β-lactamase to LANA does not appear to alter viral replication, establishment of latency, or reactivation from latency [28, 29, 32]. To confirm this reporter system works equivalently for the WT.βla and cycKO.βla viruses, we measured the frequency and expression of LANA::βla after lytic infection of mouse 3T12 fibroblasts. 3T12 cells were infected at an MOI of 10 with WT (unmarked), WT.βla, or cycKO.βla virus. At 12 hours post infection (hpi), cells were collected, and stained for β-lactamase activity using CCF2-AM, a cell-permeable β-lactamase substrate [28, 29, 32]. CCF2-AM is readily taken up by living cells, causing them to fluoresce at 520nM. If β-lactamase is present, indicating viral LANA expression, it then cleaves the substrate causing the cells to gain fluorescence emission at 448 nM. As expected, WT.βla and cycKO.βla viruses resulted in comparable frequency and expression of LANA::βla (βla^+^) following in vitro infection (Fig 1B). We next confirmed that, as reported, the β-lactamase marker did not alter reactivation phenotypes of either WT or cycKO viruses. C57BL/6 (B6) mice were infected with 1×10^6^ PFU of either WT.βla or cycKO.βla virus via intraperitoneal injection (IP). At 42 days post infection (dpi), splenocytes and peritoneal cells were collected and subjected to limiting-dilution reactivation analysis on permissive mouse embryonic fibroblasts (MEFs) as previously described [12, 33]. Briefly, latently infected splenocytes and peritoneal cells were plated on MEFs. If latent virus reactivates, the resulting virions infect and lyse the MEF monolayer. The number of latently infected cells can then be determined through nonlinear regression analysis. As previously established in comparison of WT and cycKO viruses in absence of the β lactamase fusion, the cycKO.βla virus was severely defective in reactivation from both splenocytes and peritoneal cells (Fig 1C). Taken together, these data support the previous reports that fusion of β-lactamase to LANA does not alter the biology of these viruses [28, 29, 32].

**Figure 1.**
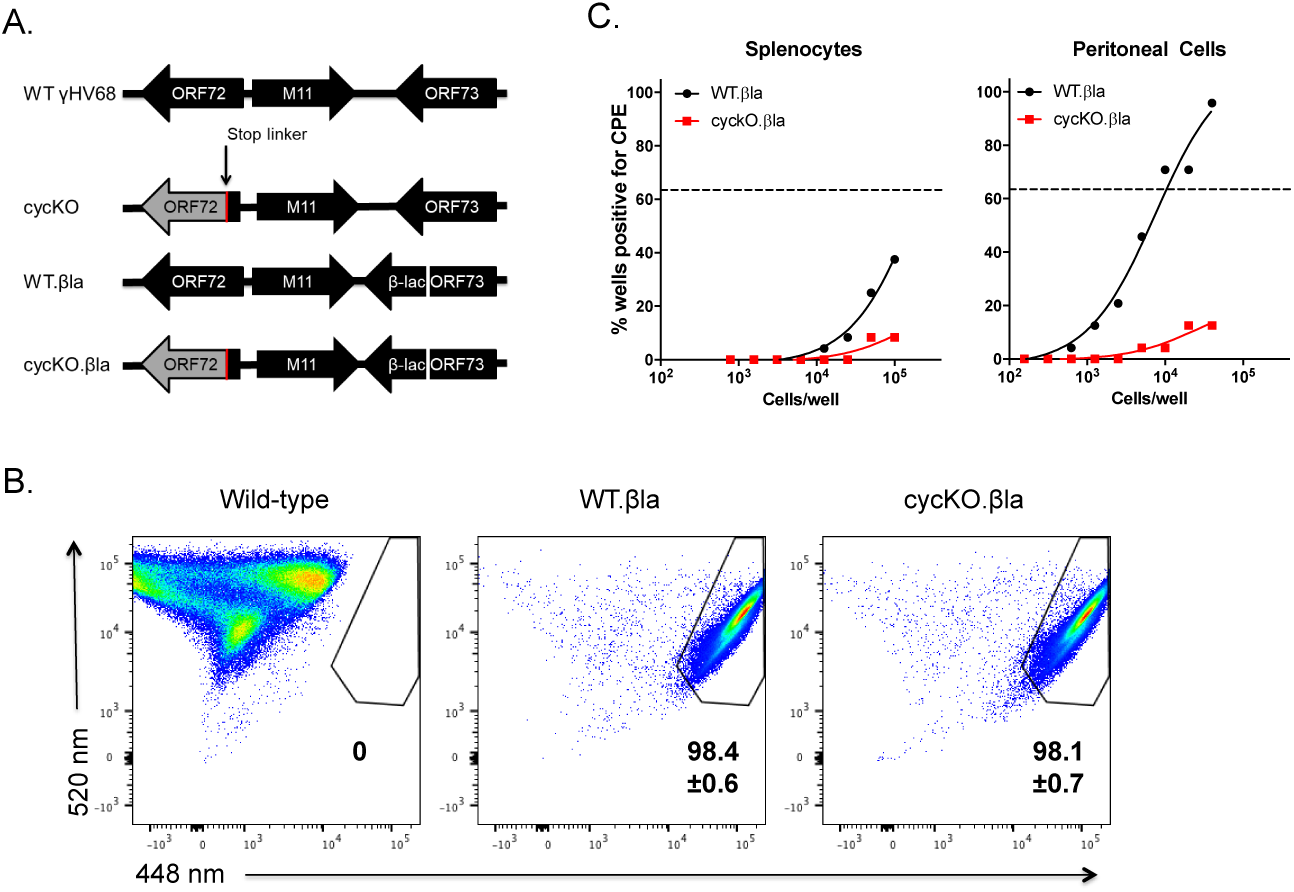
Characterization of the cyclin-deficient virus expressing a LANA β-lactamase gene fusion. (A) Schematic of wild-type virus (top), the cyclin deficient virus (second), the wild-type β-lactamase marked virus (third), and the cyclin deficient β-lactamase marked virus (bottom). Viruses described as in van Dyk 2000 and Niemeyer 2018. (B) Identification of infected cells by flow cytometry using β-lactamase (βla). 3T12 cells were infected with either wild-type unmarked γHV68, WT.βla, or cyckO.βla at an MOI of 10 pfu/cell. Cells were harvested at 12 hpi and infected cells were identified by β-lactamase activity. LANA::βla+ cells are contained within the upper right polygonal gate. Average LANA::βla+ frequencies are indicated +/-SEM n=2. (C) B6 mice were infected via IP infection with WT.βla (black) or cycKO.βla (red) viruses. At 42 dpi splenocytes (left panel) and peritoneal cells (right panel) from infected mice were plated on MEFs in a limiting-dilution fashion. Comparison of reactivation from infected cells were pooled for reactivation analysis. For each virus group 5 mice were infected and pooled for reactivation analysis.

### The cell composition of cycKO.βla infected mice is not altered compared to WT.βla infection

To determine if the cycKO virus is preferentially enriched in a particular subset of cells, we infected (B6) mice with 1×10^6^ PFU of either WT.βla or cycKO.βla via IP injection. Splenocytes were harvested at 8 dpi and 16 dpi. Eight dpi is a time point within the acute phase of infection, while 16 dpi corresponds to the establishment of latency after acute infection has been resolved [34, 35]. After collection, splenocytes were stained for LANA::βla, CD19, IgD, CD38, and CD44. These markers were used to identify B cells (CD19^+^), including germinal center B cells (CD19^+^, IgD^-^, CD38^-^) or activated B cells (CD19^+^, IgD^-^, CD44^+^). We chose to measure these populations because germinal center B cells represent an important population for γHV68 to infect and seed memory B cells [36], the primary cell type harboring long-term latent virus, and activating B cells has been show to stimulate reactivation [37]. We determined the composition of infected cells by identifying cells expressing the viral LANA::βla fusion protein (Fig 2A). We quantified germinal center B cells and activated B cells by sequentially gating on CD19^+^, IgD^-^, and CD38^-^ or CD44^+^ respectively (Fig 2B). We saw no significant differences in the expression of these markers on total or infected (βla^+^) splenocytes at 8 dpi during acute infection (Fig 2C) or at 16 dpi during latency (Fig 2D). In agreement with this, there were no differences between WT.βla or cycKO.βla virus in the frequency of βla^+^ cells that were total B cells, germinal center B cells, or activated B cells (Fig 2E). Contrary to our initial prediction, these data suggest that although the cycKO.βla virus is defective in reactivation there are no appreciable differences in the composition of the infected cells compared to WT.βla virus. Thus, there must be another explanation for the reactivation defect observed in v-cyclin deficient viruses.

**Figure 2:**
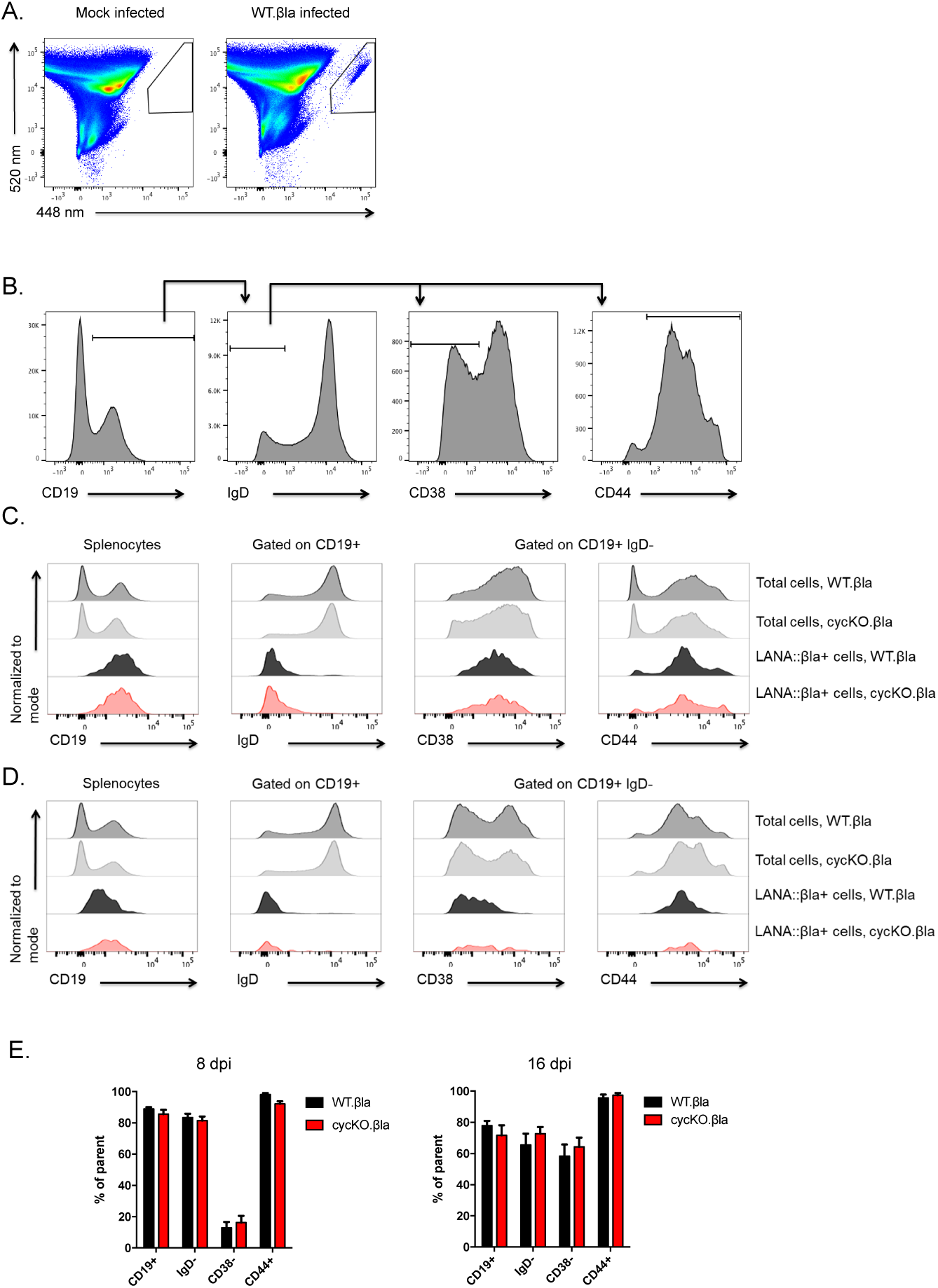
Cyclin deficient virus has a similar cellular distribution to wild-type γHV68 during primary infection of C57BL/6J mice. (A) Representative gating of LANA::βla+ splenocytes. (B) Representative gating strategy for total B cells (CD19+), germinal center B cells (CD19+, IgD-, CD38-), and activated B cells (CD19+, IgD-, CD44+). (C) and (D) representative histograms of cell surface marker expression after infection with WT.βla or cycKO.βla virus on total cells (dark grey WT.βla and light grey cycKO.βla), WT.βla infected LANA::βla+ cells (black), or cycKO.βla infected LANA::βla+ cells (red). Cells were harvested at either 8 dpi (C) or 16 dpi (D). (E) Quantification of the frequency of cells expressing cell surface markers using the gating strategy outlined in (B) with SEM shown. 8 dpi CD19, IgD and CD38 n=13. 8 DPI CD44 n=7. 16 DPI n=11. Two-tailed student t tests were performed to measure statistical significance.

### CycKO.βla virus infection results in deficient expression of viral LANA compared to WT.βla

Splenocytes, from mice infected as above, were collected at 8 and 16 dpi and analyzed by limiting-dilution nested PCR to measure the frequency of splenocytes harboring viral DNA [12, 33]. We found that there was a minor decrease in the number of cells harboring cycKO.βla virus at 8 dpi but no significant difference in the number of cells containing γHV68 DNA following infection at 16 dpi (Fig 3A). These data indicate that the reactivation defect in cycKO virus is not due to fewer cells becoming infected, consistent with previously published reports [28, 30]. However, when splenocytes were analyzed for the frequency of LANA::βla expressing cells, we found a significantly lower frequency of LANA::βla^+^ cells in mice infected with cycKO.βla (0.06%) compared to wild-type virus infected samples (0.21%) at 8 dpi. Further, this trend continued into latency with 0.03% of splenocytes at 16 dpi that were LANA::βla^+^ after WT.βla infection compared to 0.008% of splenocytes after cycKO.βla infection (Fig 3B). This difference in frequency corresponded to a decrease in the total number of LANA::βla^+^ splenocytes per mouse after infection with the cycKO.βla virus (Fig 3C). Considering an equivalent number of cells are viral DNA positive (Fig 3A), this indicates that there is a decrease in the proportion of infected cells that expressed LANA in the absence of v-cyclin. This decreased frequency of LANA::βla^+^ cells that are B cells, germinal center B cells, or memory B cells translated into a sharp decline in the number of LANA::βla^+^ cells in cycKO.βla infected mice compared to WT.βla infected mice across multiple subsets (Fig 3D). As WT.βla and cycKO.βla viruses had comparable β-lactamase expression during lytic infection of 3T12 cells (Fig 1B), these data indicate that the v-cyclin promotes the frequency of LANA expressing cells during latent infection in vivo.

**Figure 3.**
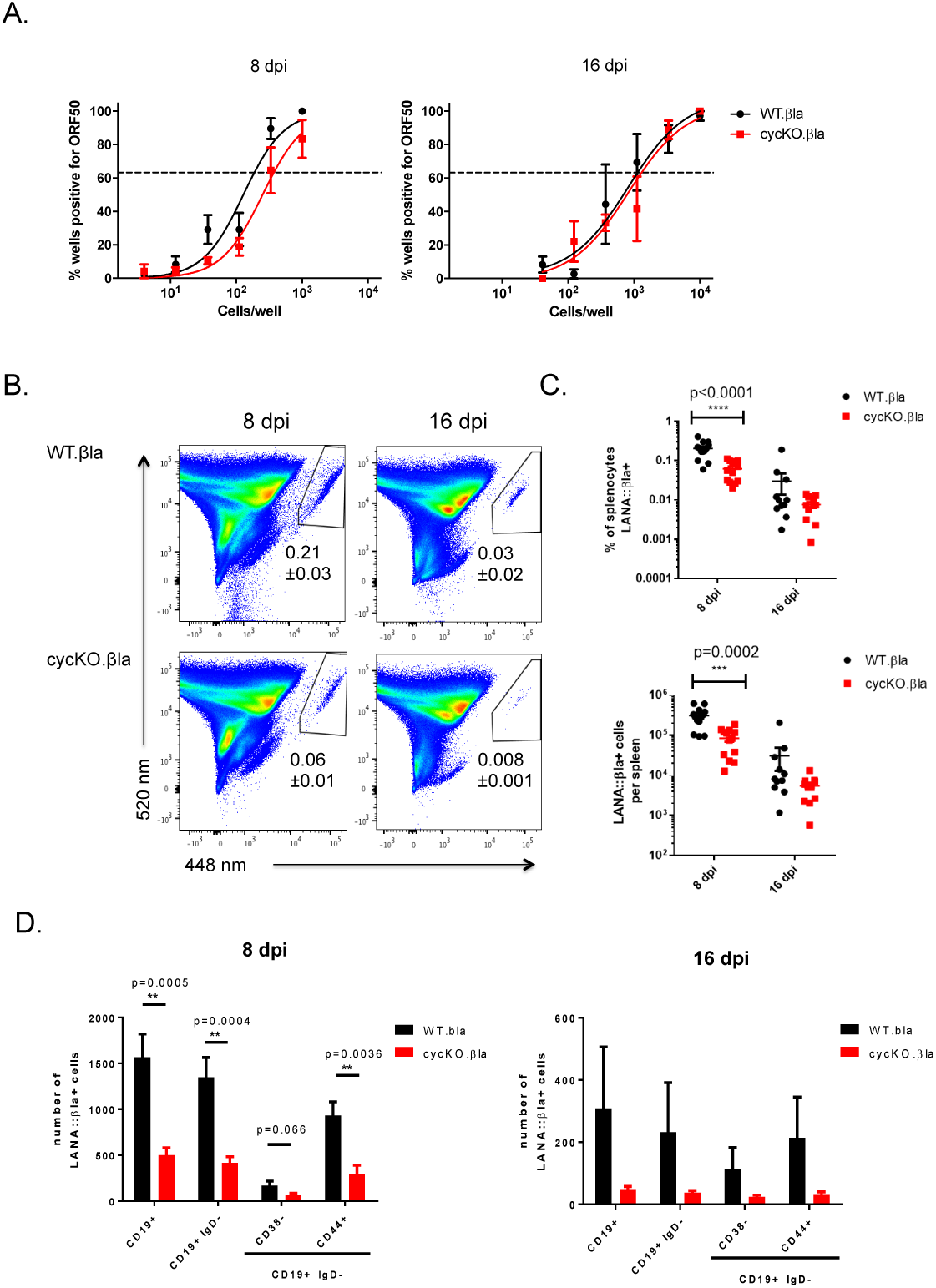
Disruption of the viral cyclin has no effect on the frequency of viral genome positive cells yet results in a reduced frequency of LANA::βla+ cells. Mice were infected via IP injection with WT.βla or cycKO.βla viruses and splenocytes were harvested at 8 or 16 dpi. (A) Limiting-dilution nested PCR of viral gene ORF 50 from WT.βla or cycKO.βla infected splenocytes. n=4 (8 dpi) or n= 3 (16 dpi) with 3-5 mice pooled per group with SEM shown. Comparisons between the LogEC(63.2) found statistical difference between the of WT.βla and cycKO.βla at 8 dpi only (p=0.009). (B) Representative pseudocolor plots identifying LANA::βla+ splenocytes, indicated in the upper right polygon. Average frequencies of LANA::βla+ cells +/-SEM is indicated below the gate. 8 dpi is shown on the left, while 16 dpi is shown on the right, with WT.βla infected mice on top and cycKO.βla infected mice on bottom. (C) Percent of LANA::βla+ cells (top) and total number of LANA::βla+ splenocytes (bottom) from each individual mouse plotted with SEM shown after infection with WT.βla (black) or cycKO.βla (red) virus. (D) Graphical representation of the number of LANA::βla+ cells expressing cell surface markers with SEM shown after infection with WT.βla (black) or cycK.βla (red) virus. Cells were stained and gated as in figure 2. 8 dpi: LANA::βla+, CD19, IgD, and CD38 n=13 and CD44 n=7. 16 dpi n= 11. Two-tailed student t tests were performed to measure statistical significance in C and D.

### The defect in LANA expression with cycKO.βla infection is observed regardless of the tissue type

While we consistently observed a decrease in the frequency of cells expressing LANA::βla after cycKO.βla infection, it remained possible that this was a tissue-specific phenotype. To address this possibility, mice were infected IP as described above and peritoneal cells were collected at 8 and 16 dpi, stained for β-lactamase, CD19, and CD5. CD19 was used to distinguish between non-B cells and B cells (CD19^+^) and CD5 expression on CD19^+^ cells was used to identify B1-a cells, which are known to harbor latent virus in the peritoneum (Fig 4A) [13, 28]. We saw no significant difference in the cellular distribution of infection between WT and cycKO viruses (Fig 4B), but a profound decrease in the frequency of LANA::βla^+^ cells in peritoneal cells harvested from cycKO.βla infected mice (Fig 4C). There was a significantly lower frequency of LANA::βla^+^ peritoneal cells after cycKO.βla infection at both 8 and 16 dpi (Fig 4D). This indicates that the v-cyclin is required for optimal LANA expression in the peritoneum and the spleen, two dominant sites for latency. Finally, to determine whether this effect was dependent on route of infection, we measured the frequency of LANA::βla^+^ cells in the lungs at 8 days post-intranasal infection (Supplemental Fig 1). Again, mice infected with cycKO.βla virus had a reduced frequency and number of LANA::βla^+^ compared to WT.βla infected mice. These data demonstrate that the v-cyclin is required for optimal LANA expression, regardless of tissue or route of infection.

**Figure 4.**
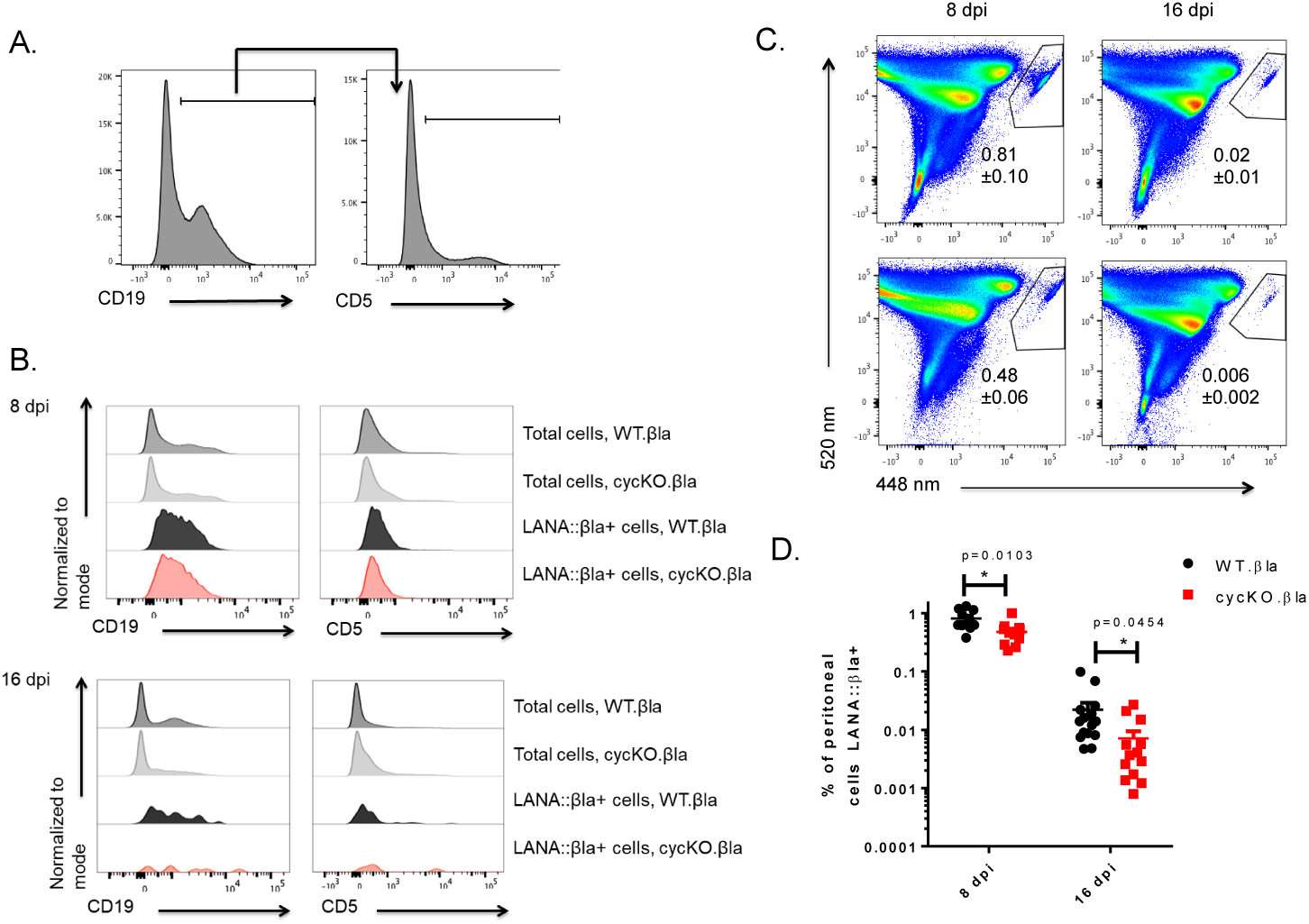
Disruption of the viral cyclin results in a reduced frequency of LANA::βla expressing cells in the peritoneum of C57BL/6J mice. Mice were infected via IP injection with WT.βla or cycKO.βla viruses and peritoneal cells were harvested at 8 or 16 dpi. (A) Representative gating strategy for peritoneal cells. (B) Representative histograms of cell surface marker expression after infection with WT.βla or cycKO.βla virus on total cells (dark grey WT.βla and light grey cycKO.βla), WT.βla βla+cells (black), or cycKO.βla βla+ cells (red) at 8 dpi (top panel) and 16 dpi (bottom panel). (C) Representative pseudocolor plots identifying βla+ cells in the upper right polygon at 8 dpi (left panel) or 16 dpi (right panel) after WT.βla (top) or cycKO.βla (bottom) virus infection. The frequency of peritoneal cells that are βla+ is indicated below the gate +/-SEM. (D) Percent of cells that are βla+ with +/-SEM shown after infection with WT.βla (black) or cycKO.βla (red) virus. 8 dpi: WT.βla n=9 and cycKO.βla n=10. 16 dpi: n=11. Two-tailed student t tests were performed to identify statistical significance.

### The v-cyclin promotes the frequency of LANA expressing cells in immunodeficient, CD8-deficient mice

The v-cyclin is required for optimal reactivation across both immunocompetent and immunodeficient genetic backgrounds [30, 33, 38]. CD8-deficient (CD8^-/-^) mice, which lack CD8 T cells, have a significant increase in the number of latently infected cells relative to B6 controls [39]. Despite the overall increase in the number of latently infected cells, the cycKO.βla virus is still defective in reactivation in these mice [33]. We therefore tested whether the v-cyclin was required to promote LANA expression in CD8^-/-^ mice.

CD8-/-mice were infected via IP inoculation with either WT.βla or cycKO.βla virus. Splenocytes were harvested at 16 dpi and stained for β-lactamase activity, CD19 expression, and IgD and CD38 expression on CD19^+^ cells. We found that, as with B6 mice, there were no differences in cellular distribution of LANA::βla^+^ between WT and cycKO viruses (Fig 5A, 5C).

**Figure 5.**
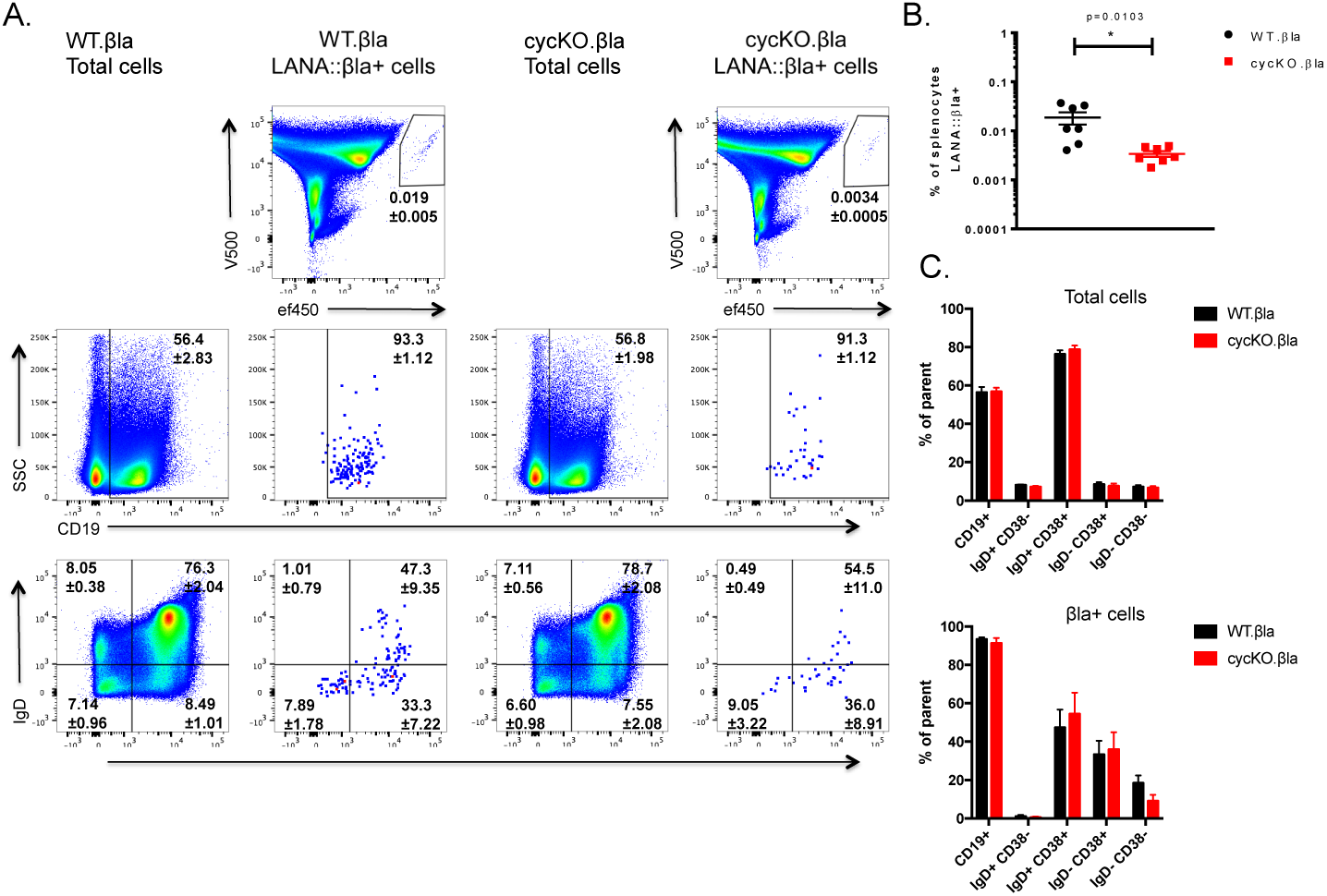
Disruption of the viral cyclin results in a reduced frequency of LANA::βla expressing cells in the spleen of CD8-/-mice. At 16 dpi, splenocytes were collected and stained for β-lactamase activity and cell surface markers CD19, CD38, and IgD. CD38 and IgD samples have been previously gated on CD19+ cells. (A) Representative pseudocolor plots are shown identifying LANA::βla+ cells (top row). LANA::βla+ cells are found within the upper right polygon with the average percent of cells expressing LANA::βla +/-SEM indicated below the gate. Expression of CD19, IgD, and CD38 on total cells after WT.βla virus or cycKO.βla virus infection is shown in the indicated columns. Expression of CD19, IgD, and CD38 on LANA::βla+ cells is shown in the indicated columns. The average percent of cells that fall within each gate is indicated +/-SEM. (B) Graphical representation of the percent of cells that are LANA::βla+ after WT.βla (black) or cycKO.βla (red) infection with SEM. (C) Graphed are the average percent of cells that are CD19+ and the percent of CD19+ cells that are IgD+/CD38-, IgD+/CD38+, IgD-/CD38+, and IgD-/CD38-after WT.βla (black) or cycKO.βla (red) infection. Total cells are graphed on the top while LANA::βla+ gated cells are shown on the bottom, both with SEM plotted. Two experiments were performed with 3-4 WT.βla and cycKO.βla infected mice per experiment. WT.βla: n=7; cycKO.βla: n=6

Importantly, the defect in LANA::βla expression in cycKO infected splenocytes is still maintained, with a 5.6-fold decrease in the frequency of splenocytes that are βla+ after cycKO.βla infection (0.0034%) compared to WT.βla infection (0.019%) (Fig 5B). We also analyzed peritoneal cell infection at 16 dpi. Peritoneal cells from mice infected as above were collected and stained for β-lactamase activity, CD19, B220, and CD5. The cycKO.βla defect was also present in the peritoneal compartment, with only 0.054% of peritoneal cells LANA::βla^+^ in cycKO infected samples compared to 0.496% LANA::βla^+^ cells after WT.βla infection (Fig 6A, 6B). Interestingly, we detected a modest shift in the peritoneal composition of LANA::βla^+^ cycKO.βla infected cells: 25% of cycKO.βla infected LANA::βla^+^ cells were CD19^+^ compared to 12% of WT.βla infected LANA::βla^+^ cells (Fig 6C). This difference mirrors a change in the total frequency of CD19^+^ cells in the peritoneum after cycKO.βla infection (Fig 6C). When analyzing the composition of infected B cells by B220 and CD5 expression, the LANA::βla^+^ cells were found in B1-a, B1-b, and B2 cells, with a higher prevalence in B1 populations. Of WT.βla infected LANA::βla^+^ cells: 4% were B2 cells, 5% were B1-a cells, and 8% were B1-b cells. Of the cycKO.βla infected LANA::βla+ cells: 10% were B2 cells, 8% were B1-a cells, and 10% were B1-b cells (Fig 6C). Interestingly, we have previously identified a similar trend in p18Ink4c deficient mice, a mouse strain in which there is an overall increase in reactivation [28], similar to the CD8-/-mice.

**Figure 6.**
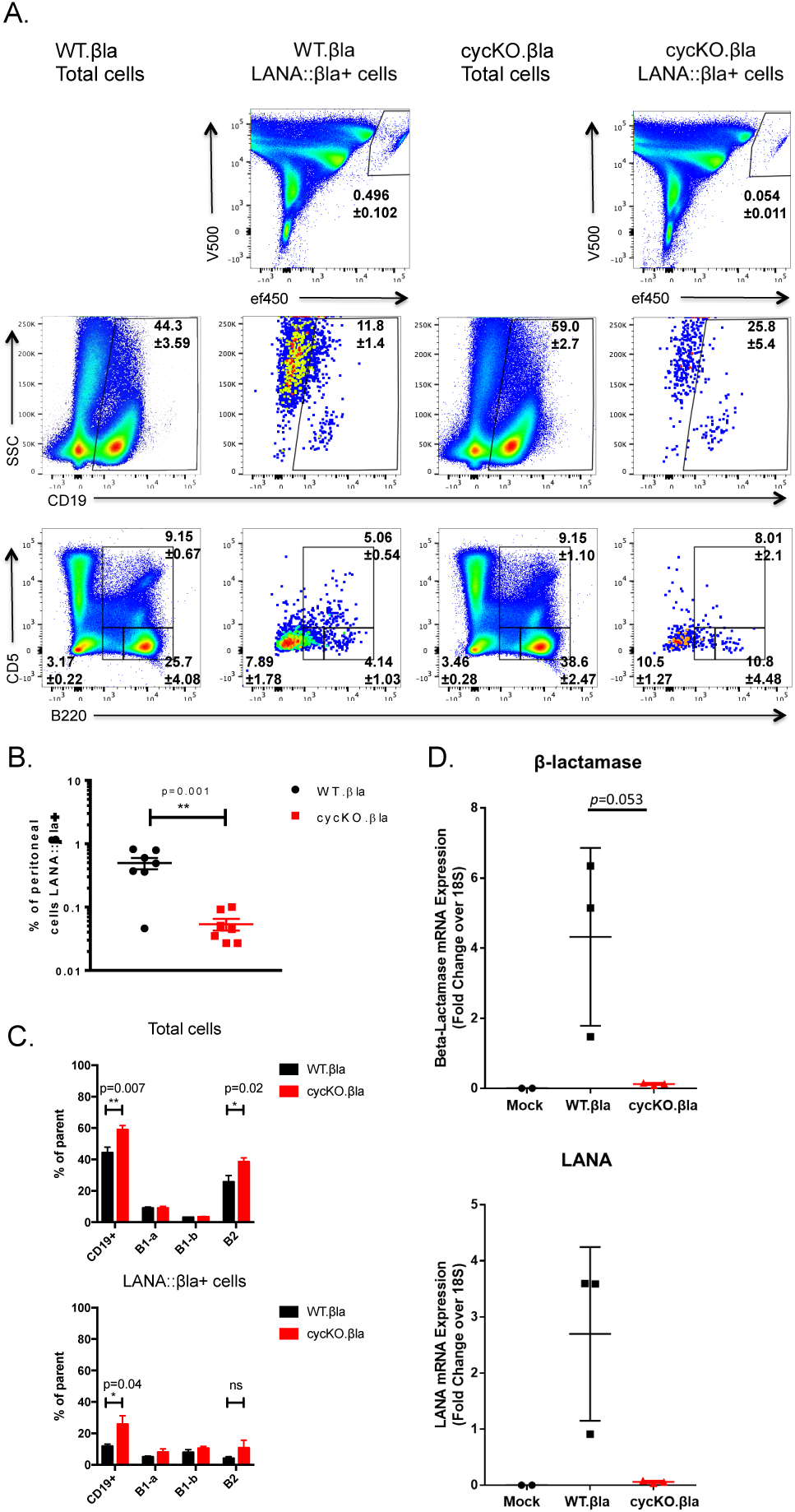
Disruption of the viral cyclin results in a reduced frequency of LANA::βla expressing cells in the peritoneum of CD8-/-mice. CD8-/-mice were inoculated with WT.βla virus or cycKO.βla virus via IP injection. At 16 dpi, peritoneal cells were collected and stained for β-lactamase activity and cell surface markers CD19, B220, and CD5. (A) Representative pseudocolor plots are shown identifying LANA::βla+ cells (top row). LANA::βla+ cells are found within the upper right polygon with the average percent of cells expressing LANA::βla +/-SEM indicated below the gate. Expression of CD19, B220, and CD5 on total cells after WT.βla virus or cycKO.βla virus infection is shown in the indicated columns. Expression of CD19, B220, and CD5 on LANA::βla+ cells is shown in the indicated columns. The average percent of cells that fall within each gate is indicated +/-SEM. (B) Graphical representation of the percent of cells that are LANA::βla+ after WT.βla (black) or cycKO.βla (red) infection with SEM. (C) Graphed are the average percent of cells that are CD19+, B1-a (CD5+), B1-b (B220 intermediate), and B2 (B220 high) after WT.βla (black) or cycKO.βla (red) infection. Total cells are graphed on the top while LANA::βla+ gated cells are shown on the bottom, both with SEM plotted. Two experiments were performed with 3-4 WT.βla and cycKO.βla infected mice per experiment. WT.βla: n=6 cycKO.βla: n=7. (D) Relative expression of β-lactamase (top) and LANA (bottom). RNA was isolated from infected peritoneal cells and subjected to quantitative RT-PCR analysis using primers directed against β-lactamase (top) and LANA (bottom). mRNA expression levels depicted were normalized to 18S levels, with differences between WT. βla and cycKO.βla as noted.

To determine LANA::βla gene expression independent of enzymatic activity, we isolated WT or cycKO infected peritoneal cells and measured both β-lactamase and LANA RNA by qRT-PCR (Fig 6D). Similar to analysis by enzymatic activity, these data demonstrate a difference in latent gene expression at the RNA level between WT and cycKO infected cells at 16 dpi. These data indicate that the v-cyclin is required for optimal LANA gene expression in multiple tissues and in both immunocompetent and immunodeficient hosts.

### The v-cyclin is dispensable for reactivation in LANA expressing cells

The reduced frequency of LANA expressing cells after cycKO.βla infection in both B6 and CD8^-/-^ mice correlates with the reactivation defect of the cycKO.βla virus observed in these mice. It has also been previously reported that sort purifying LANA::βla^+^ cells can enrich for cells capable of reactivation [29]. Based on these observations, we postulated that the defect in reactivation of cycKO viruses may be a direct consequence of the reduced frequency of LANA expressing cells. To this end, we used flow sorting to purify LANA::βla^+^ cells from mice infected with either WT.βla and cycKO.βla virus and measured reactivation capacity ex vivo. Given the low frequency of βla^+^ cells in healthy B6 mice (Fig 3B, 4C, and 5A), we sorted LANA::βla^+^ cells from CD8^-/-^ mice. CD8-/-mice were infected with WT.βla or cycKO.βla virus via IP injection with peritoneal cells harvested at 16 dpi, stained for β-lactamase expression and FACS purified into either LANA::βla^+^ or LANA::βla^-^ populations (Fig 7A), followed by flow cytometric analysis of sort purity (Fig 7B). For each sort, we recovered ∼25,000-100,000 WT.βla infected LANA::βla^+^ cells, ∼21,000-24,000 cycKO.βla infected LANA::βla^+^ cells, and ∼1.5-2×10^6^ LANA::βla^-^ cells for each virus. Bulk, LANA::βla^+^, and LANA::βla^-^ cells were plated by serial dilution on MEF monolayers and assessed for reactivation 21 days post-plating. As expected, in the pre-sorted population the cycKO.βla virus showed a reactivation defect relative to WT.βla infected peritoneal cells (Fig 7C). Reactivation in the LANA::βla^-^ population was extremely low. In contrast to the low frequencies of reactivation present in the LANA::βla^-^ population, LANA::βla^+^ cultures had much higher frequencies of reactivation (Fig 7C). Notably, when the number of LANA expressing cells analyzed for reactivation was normalized between WT.βla and cycKO.βla infected samples, the cycKO.βla had equivalent reactivation to WT.βla virus (Fig 7C). These data directly demonstrate that LANA expressing cells are enriched in their ability to reactivate from latency, and provide direct evidence that a primary function of the v-cyclin is to promote the frequency of LANA expressing cells during latent infection in vivo. They further indicate that the defect in reactivation observed in bulk cultures from cycKO infected mice can be directly attributed to a reduced frequency of LANA expressing cells in the cycKO infected latent reservoir.

**Figure 7.**
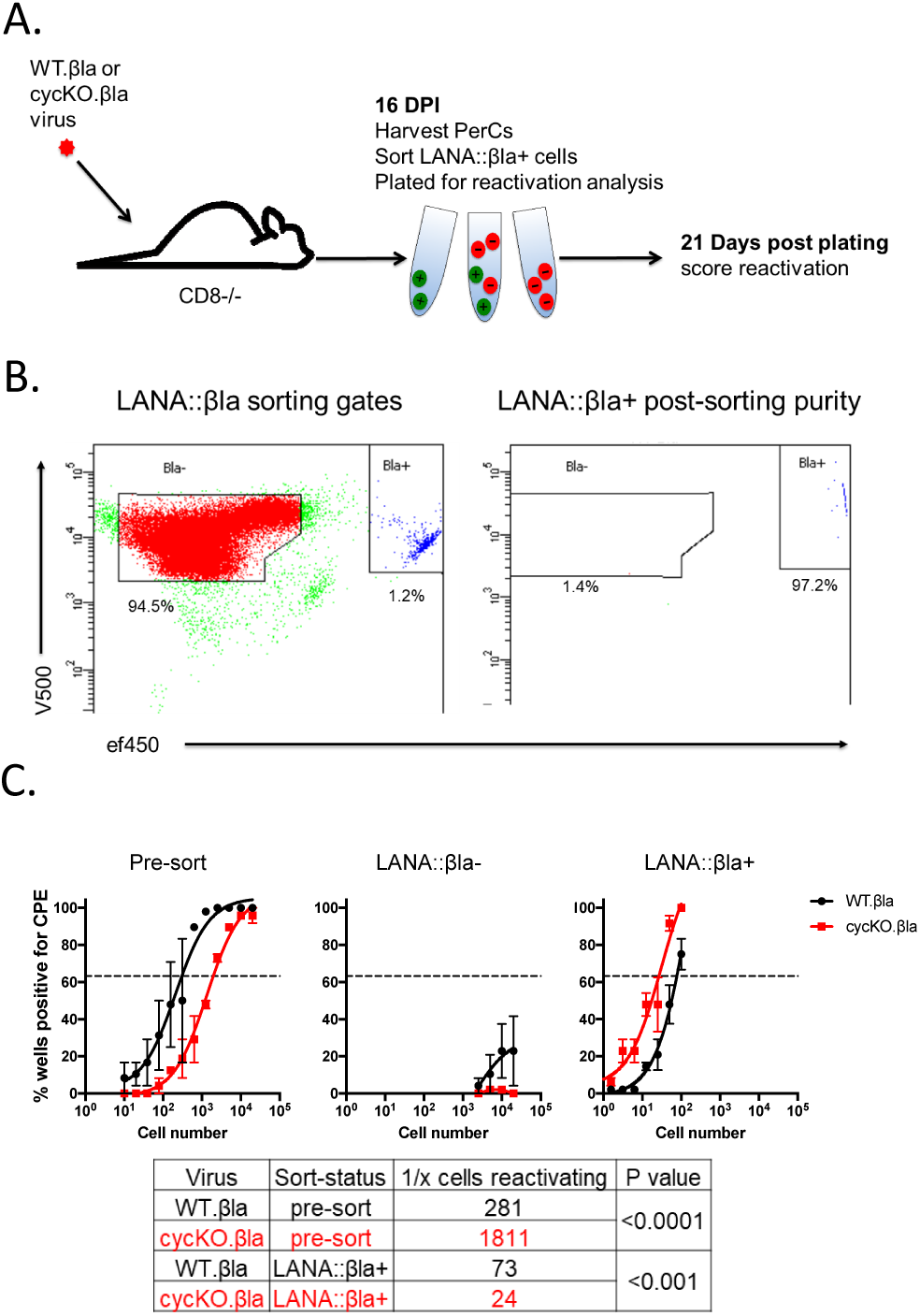
The viral cyclin is dispensable for reactivation in latently infected cells that express LANA::βla. (A) Experiment schematic. CD8-/-mice were infected via IP injection with WT.βla or cycKO.βla viruses and peritoneal cells were harvested at 16 dpi. Cells were stained for β-lactamase activity and LANA::βla+ cells were sorted by FACS. (B) Gating strategy of LANA::βla sort (left) and purity of post-sort WT.βla infected LANA::βla+ cells (right). LANA::βla+ cells are located in the upper right polygon and LANA::βla-cells are located in the upper left polygon. The percent of events within each gate is indicated below the gate. (C) WT.βla (black) and cycKO.βla (red) infected pre-sorted (left), LANA::βla-(middle), and LANA::βla+ (right) cells were subjected to limiting-dilution reactivation analysis. Reactivation was measured 21 days after plating sorted and pre-sorted cells on permissive MEFs. Linear regression with comparison of the LogEC(63.2) found that there was a statistical difference in reactivation between LANA::βla+ WT.βla and cycKO.βla infected cells and pre-sorted WT.βla and cycKO.βla infected cells (p<0.0001). n=2 independent experiments with a total of 15 WT.βla infected mice and 30 cycKO.βla infected mice. The table below lists the number of cells plated to reach CPE in 63.2% (the dotted line) of the wells plated for reactivation, corresponding to the number of cell required to find at least 1 reactivating cell.

## Discussion

The balance between latency and reactivation is of critical importance in γHV infection and disease progression. Chronic infection with γHV through maintenance of latency and reactivation has long been associated with virus-induced malignancies [1]. Here, we find that a primary function of the v-cyclin is to promote latent gene expression (i.e. LANA), to create a reactivation competent latent reservoir. Our findings suggest that γHV68 latency is not a uniform state. Indeed, it appears that some latently infected cells are more prone to reactivation while others seem refractory. In the work presented here, we show that LANA expression is a strong correlate with reactivation capacity, while cells that fail to express LANA have limited reactivation potential (Fig 7C). Therefore, we propose a more comprehensive model of gammaherpesvirus reactivation where either the viral cyclin, or potentially host cyclins, promote reactivation by altering the state of the latently infected cell (Fig 8). In this specific instance, expression of v-cyclin increases the pool of LANA expressing latently infected cells, which are more permissive to reactivation from latency (Fig 8).

**Figure 8.**
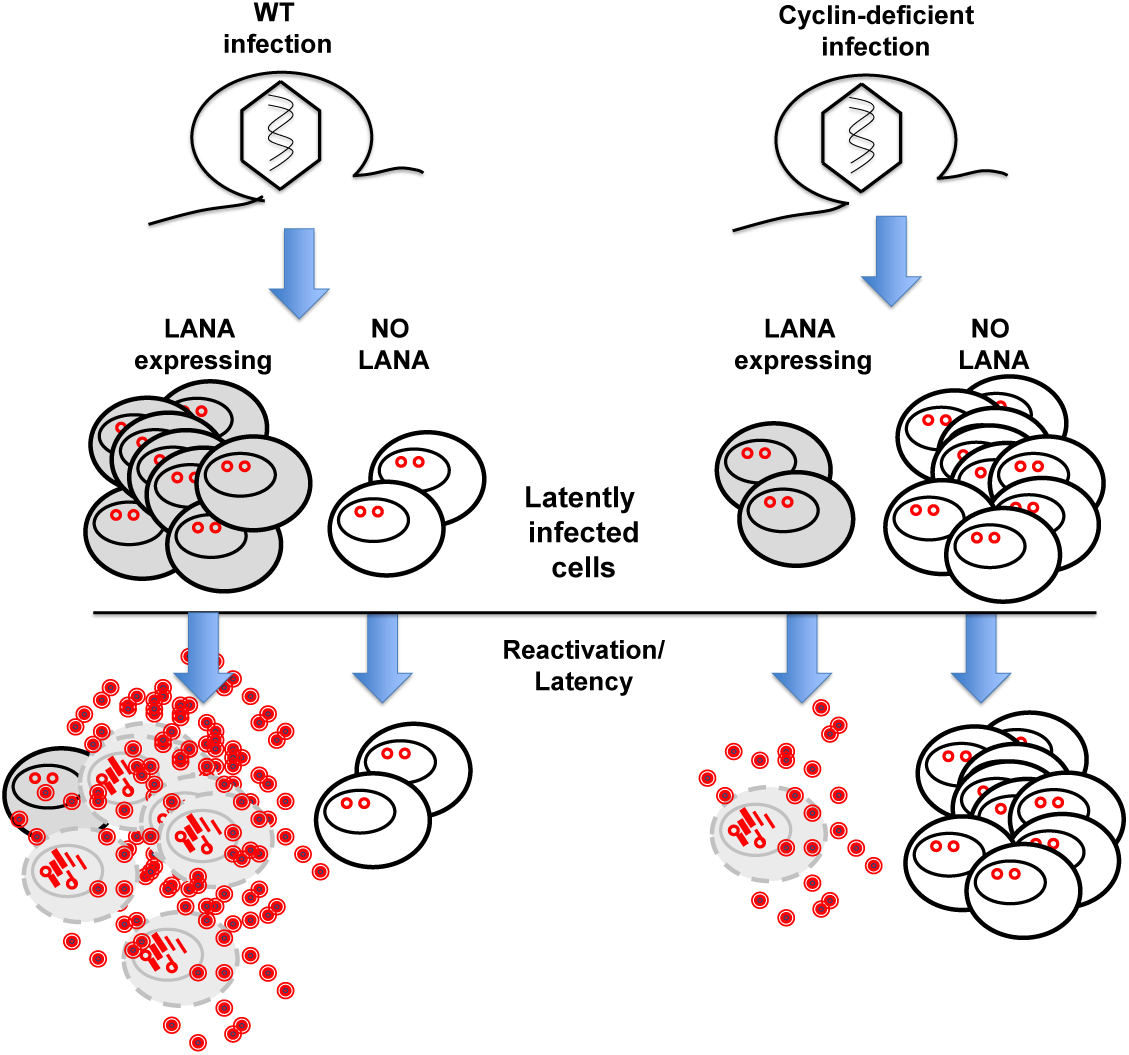
The viral cyclin drives reactivation through increasing the pool of LANA expressing, reactivation-competent, infected cells. Wild-type γHV68 infection results in establishment of latency (depicted by nuclear viral episomes). At least two distinct populations of latently infected cells arise, cells expressing viral LANA (gray) and those lacking detectable LANA expression (white). The LANA expressing cells are permissive to reactivation and will readily reactivate when triggered. Latent cells lacking LANA are incapable of reactivation, and instead remain dormant in latency. Infection with a viral cyclin-deficient γHV68 virus also results in establishment of latency, with equivalent numbers to wild-type infection. However, without the viral cyclin, latency is skewed to reactivation incompetent, LANA negative cells. The cyclin-deficient infected cells which do express LANA are still able to reactivate, as efficiently as wild-type, but there are diminished numbers of these cells ultimately leading to the reactivation defect.

The notion that viral latency is a diverse and complex state of infection was originally defined in EBV infection, in which there are several distinct types of latency [40]. These diverse latency stages are distinguished by variable expression of viral genes in a manner that is mimicked with cyclin-deficient γHV68 infection. Further, different EBV latency programs are associated with specific clinical outcomes and pathologies [41, 42]. While our findings likely reflect a conserved feature of biology amongst gammaherpesvirus, similar trends can be observed in latent infection across virus families. For example, in HIV infection, virus persists within the host through latency in many cells types, including CD4 T cell subsets and myeloid cells [43, 44]. Since these reservoirs cannot be cleared by therapeutics or the host immune system, one potential strategy for eradication of the virus is to trigger reactivation, resulting in death of the cell by the virus or the host immune system [45]. One barrier to this approach is the fact that distinct populations of latently infected cells appeared differentially responsive to reactivation stimuli [46].

One important question raised by our findings is how broadly the v-cyclin is required for latent gene expression: is this effect specific to LANA, or does it apply more broadly to additional viral genes (e.g. M2)? In terms of how the v-cyclin is regulating latent gene expression, it is possible that loss of v-cyclin is associated with a defect in epigenetic remodeling, resulting in broad repression of viral gene expression. While the v-cyclin promotes the frequency of LANA expressing cells, it is notable that the LANA expression during latency can occur independent of the v-cyclin. In our previous studies, we have identified two contexts in which reactivation can occur in a v-cyclin independent manner: i) genetic, or physiological, loss of p18Ink4c enables robust reactivation of γHV68 in cycKO infection [27, 28], and ii) host cyclin D3 is capable of fulfilling the role of v-cyclin in driving reactivation in a cycKO background, albeit with a decreased efficiency [47]. Based on these observations, it is possible that cycKO infected cells with LANA expression may reflect cells with either increased cellular D type cyclin expression and/or decreased p18Ink4c expression, allowing both LANA expression and reactivation. Although our data emphasize that the v-cyclin promotes LANA expression and that LANA expressing cells are reactivation competent, it remains to be tested whether the v-cyclin supports additional features for optimal reactivation capacity.

The work described here documents a critical link between v-cyclin and viral LANA expression in reactivation from latency. Further, our findings strongly suggest that the latently infected reservoir is diverse in gene expression and reactivation capacity. These data identify a v-cyclin/LANA axis that is critical for reactivation from latency and emphasize that efforts to manipulate this axis may require a combinatorial approach that targets both v-cyclin dependent and independent processes to effectively disrupt the latent reservoir.

## Methods and Materials

### Cell lines and viruses

3T12 mouse fibroblast cells (ATCC CCL-164) were cultured in 5% FBS/DMEM with 20 units of penicillin and 20 μg of streptomycin per mL and 4 mM L-glutamine. MEFs were isolated as described and cultured in 10% FBS/DMEM with 20 units of penicillin per mL, 20 μg of streptomycin per mL, 4 mM L-glutamine, and fungizone at 250 ng/mL [48]. Generation of the WT.βla and cycKO.βla viruses has been previously described [28, 29, 32].

### Mice

C57BL/6 (B6) mice were obtained from the Jackson Laboratory (Stock # 000664). CD8α-/-mice on the B6 background (CD8-/-) were obtained from the Jackson Laboratory (Stock # 002665) and have been previously described [39]. CD8-/-mice were bred in house at the University of Colorado Denver Anschutz Medical Campus in accordance with University regulations and Institutional Animal Care and Use Committee.

### Flow cytometry analysis

Spleens were collected and splenocytes were isolated in a single cell suspension after being passed through a 100 micron filter. Splenocytes were then subjected to red blood cell lysis by treatment with red blood cell lysis buffer (Sigma # R7757) per manufacturer’s recommendation. Peritoneal cells were collected with 10 mLs of cold 1% FBS DMEM. β-lactamase activity was detected using the LiveBLAzer FRET-BG/Loading Kit with CCF2-AM (ThermoFischer Scientific # K1025) as previously described [17, 29, 32]. Cell surface antibodies used were CD19-AlexaFluor 700 (clone eBio1D3, eBioscience # 56-0193-81), CD38-APC (clone 90,eBiosciences #17-0382-81), IgD-APC-Cy7 (clone 11-26c.2a, Biolegend # 405716), and CD5-APC (clone 53-7.3, eBioscience # 17-0051-81). Fc blocking antibody 24G2 was used in staining to prevent antibody binding to cellular Fc receptors.

### Limiting-dilution analysis

Mice were inoculated with either WT.βla or cycKO.βla at 1×10^6^ PFU/mouse via IP injection. After 8 and 16 days, splenocytes and peritoneal cells were collected as above and analyzed by either flow cytometry or plated for reactivation or PCR analysis. By Poisson distribution, the number of cells plated corresponding to 63.2% of the wells positive is the frequency at which there is at least one reactivating or genome positive cell, respectively.

#### Reactivation analysis

Cells were subjected to serial limiting dilution analysis, and plated on highly permissive MEF monolayers for quantification of virus cytopathic effect as previously described [12, 33]. To control for any preformed virus, mechanically disrupted peritoneal cells were plated in parallel; no monolayer disruption was observed in disrupted cells.

#### LD-PCR analysis

Cell dilutions were subjected to in-plate DNA isolation and nested-PCR for single copy sensitivity detection of viral gene 50 DNA, with plasmid sensitivity controls included on each plate, as previously described [12,33].

### Quantitative-PCR analysis

CD8-/-mice were infected with 1×10^6^ PFU of either WT.βla (n=3) or cycKO.βla (n=3) virus or mock infected (n=2) via IP injection. At 16 dpi, peritoneal cells from individual mice were harvested from each mouse, pelleted at 1,000xg for 10 min, resuspended in RLT buffer containing β-mercaptoethanol and then frozen at −80°C. Cells were then thawed and homogenized via Qiashredder columns and RNA was isolated using the RNeasy Micro Kit. DNA was removed from the samples by treating with Turbo DNase as per the manufacturer’s recommendations (ThermoFisher). cDNA was synthesized using Superscript II Reverse Transcriptase (ThermoFisher). Primers for SYBR Green qPCR were designed using Primer3. Primers used were: LANA Forward 5’-ATCAGGGAATGCGAAGACAC, LANA Reverse 5’-GTGCCTGGTACCAAGGGTAA, β-lactamase Forward 5’-GCTATGTGGCGCGGTATTAT, β-lactamase Reverse 5’-AAGTTGGCCGCAGTGTTATC. iQ SYBR Green Supermix was used for the qPCR reactions (Bio-Rad) and qPCR was performed with technical triplicates from the peritoneal cell cDNA of each mouse, and run on the QuantStudio 7 Flex instrument.

### FACS sorted reactivation

CD8-/-mice were infected with 1×10^6^ PFU of either WT.βla or cycKO.βla virus via IP injection. At 16 dpi, peritoneal cells were collected and combined for each virus group. For each virus group, 1×10^6^ cells were set aside as “pre-sorted” cells. The remaining cells were stained for β-lactamase then washed and resuspended in 2% FBS in PBS. These cells were then sorted by the Clinical Immunology Flow Core with the University of Colorado Anschutz Medical Campus. Cells were gated as single cells and then sorted into βla^+^ or βla^-^ populations. A small number of LANA::βla^+^ WT.βla infected cells were tested for purity after the sort had concluded. The purity of the LANA::βla^+^ cells was measured in the WT.βla infected samples and found to be 97.3% pure. A corresponding purity check was not performed for the cycKO.βla infected samples due to a lower total number of cells recovered. Pre-sorted, LANA::βla^+^, or LANA::βla^-^ cells were diluted into 10% FBS in DMEM and plated onto permissive MEFs in a limiting-dilution fashion as previously described [12, 33]. Pre-sorted and LANA::βla^-^ cells were plated at starting concentrations of 2×10^4^ cells per well while LANA::βla^+^ cells were plated at a starting concentration of 100 cells per well. Three weeks after plating cells, reactivation was measured by observation of cytopathic effect on the MEF cells.

### Statistical analysis and software

Flow cytometric analysis was performed using FlowJo V.10.0.8r1. Graphs were generated and statistical analysis were performed using GraphPad Prism 7.0a. Limiting-dilution curves were created by performing a non-linear regression, log(agonist) vs. response-using the “EC anything” regression equation where F was set to 63.2, with top and bottom of the curves constrained to 100 and 0 respectively. Comparisons of the LogECF were used to determine statistical significance. Unpaired student t-tests were performed as mentioned. Quantitative-PCR data was analyzed using the Pfaffl method [49] and graphed using GraphPad Prism 7.

