## Supplemental figure 1 legend for "The gammaherpesvirus 68 viral cyclin facilitates reactivation by promoting latent gene expression"

**Supplemental Figure 1. Disruption of the viral cyclin results in a reduced frequency of LANA::βla+ cells in the lungs of C57BL/6J mice after intranasal infection** Mice were infected via I.N. inoculation with WT.βla or cycKO.βla viruses and lung were harvested at 8 dpi. (A) Representative pseudocolor plots identifying βla+ lung cells in the upper right polygon. Frequency of βla+ cells is indicated below the gate +/- SEM. (B) The percent of lung cells that are βla+ for each mouse is plotted with SEM shown after infection with WT.βla (black) or cycKO.βla (red). (C) The total number of βla+ cells per lung for each mouse is plotted with SEM shown after infection with WT.βla (black) or cycKO.βla (red). WT.βla n=6 cycKO.βla n=7. Two-tailed student t test was used for statistical analysis.
