## Supplemental figure 1 for "The gammaherpesvirus 68 viral cyclin facilitates reactivation by promoting latent gene expression"

### Supplemental figure 1: Disruption of the viral cyclin results in a reduced frequency of LANA::βla+ cells in the lungs of C57BL/6J mice after intranasal infection

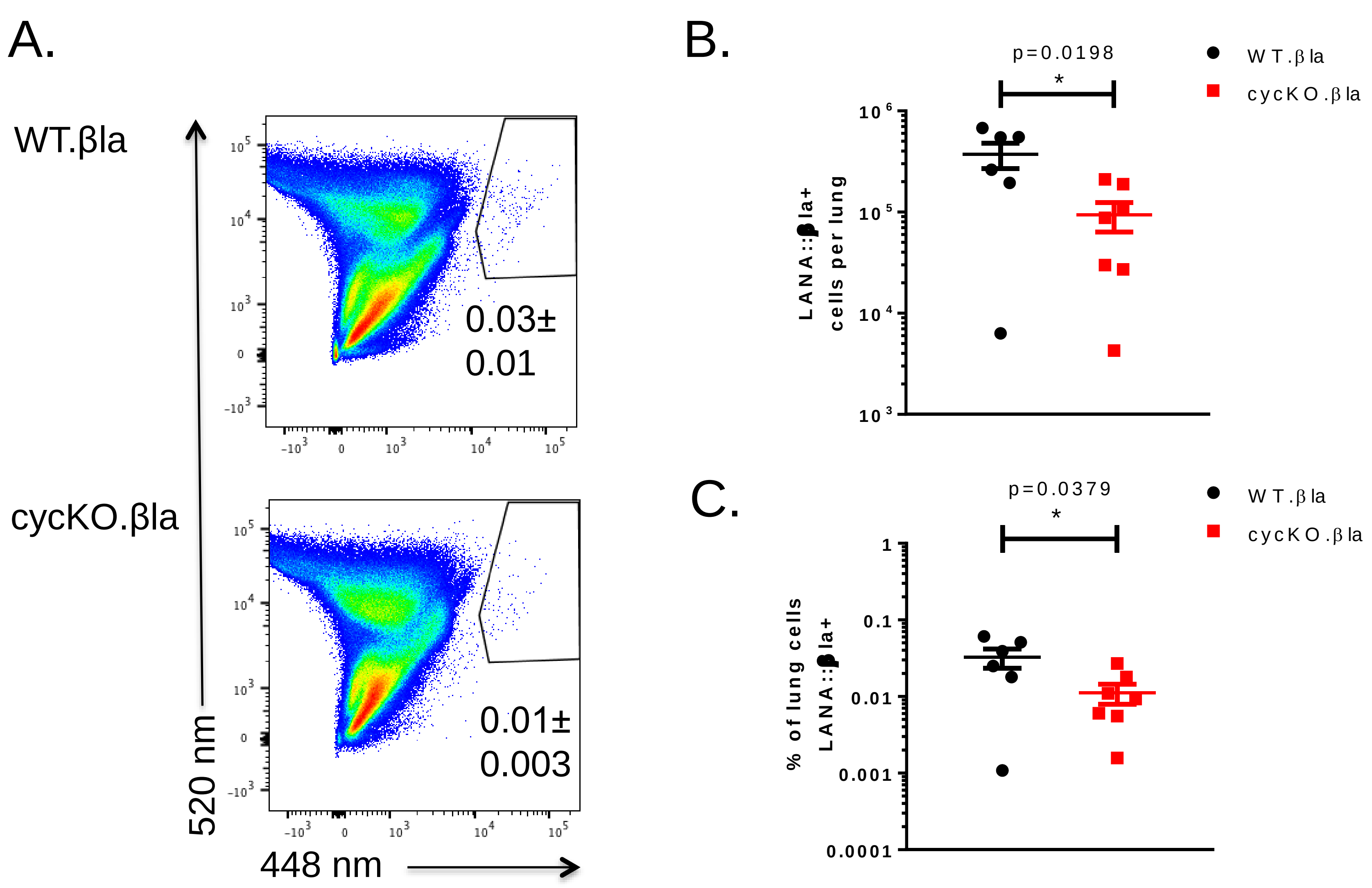
